# The acid ceramidase/ceramide axis controls parasitemia in *Plasmodium yoelii*-infected mice by regulating erythropoiesis

**DOI:** 10.1101/2022.03.08.483432

**Authors:** Anne Günther, Matthias Hose, Fabian Schumacher, Burkhard Kleuser, Kai Matuschewski, Karl S. Lang, Erich Gulbins, Jan Buer, Astrid M. Westendorf, Wiebke Hansen

**Author notes:** Corresponding author: Prof. Dr. Wiebke Hansen, Institute of Medical Microbiology, University Hospital Essen, University Duisburg-Essen, Hufelandstr. 55, 45147 Essen, Germany.

## Abstract

Acid ceramidase (Ac) is part of the sphingolipid metabolism and responsible for the degradation of ceramide. As bioactive molecule, ceramide is involved in the regulation of many cellular processes. However, the impact of cell-intrinsic Ac activity and ceramide on the course of *Plasmodium* infection remains elusive. Here, we use Ac-deficient mice with ubiquitously increased ceramide levels to elucidate the role of endogenous Ac activity in a murine malaria model. Interestingly, ablation of Ac leads to alleviated parasitemia associated with decreased T cell responses in the early phase of *Plasmodium yoelii* (*P. yoelii*) infection. Mechanistically, we identified dysregulated erythropoiesis with reduced numbers of reticulocytes, the preferred host cells of *P. yoelii*, in Ac-deficient mice. Furthermore, we demonstrate that administration of the Ac inhibitor carmofur to wild type mice has similar effects on *P. yoelii* infection and erythropoiesis. Notably, therapeutic carmofur treatment after manifestation of *P. yoelii* infection is efficient in reducing parasitemia. Hence, our results provide evidence for the involvement of Ac and ceramide in controlling *P. yoelii* infection by regulating red blood cell development.

## Introduction

Malaria remains one of the most life threatening infectious diseases in the world, with estimated 241 million cases and 627,000 related deaths in 2020 (1). Malaria is caused by infections with the parasite *Plasmodium* that undergoes a complex life cycle within its hosts. In the mammalian host, plasmodia invade red blood cells (RBCs) for their asexual propagation. Depending on the species, the parasites favour different RBC maturation stages for their intraerythrocytic development. While the most lethal human malaria species, *P. falciparum* is able to infect both, mature erythrocytes and immature reticulocytes, the most widespread parasite *P. vivax* is restricted to invade the latter (2-4). The formation of RBCs, known as erythropoiesis, primarily takes place in the bone marrow, where hematopoietic stem cells give rise to multipotent progenitors that stepwise differentiate into erythroblasts and reticulocytes (5). Early reticulocytes egress from the bone marrow and constitute 1-3% of circulating RBCs, before they develop into mature erythrocytes (6). During erythropoiesis, the cells start to produce haemoglobin, lose nuclei and organelles, and induce plasma membrane remodelling. While the erythroid lineage marker Ter119 is induced in erythroblasts and subsequently persistently expressed, expression of the transferrin receptor 1 (CD71) gets lost during the transition from reticulocytes to mature erythrocytes (7).

During the erythrocytic stage of malaria, CD4^+^ T cells play a crucial role in regulating the immune response. By secreting IFN-γ, they activate other immune cells, e.g. macrophages, which are necessary for parasite clearance and control. Moreover, CD4^+^ T cells provide help for B cells to induce protective responses against the blood-stage parasites (8). Recently, we showed that modulations of the sphingolipid (SL) metabolism enhance T cell activity during *P. yoelii* infection (9). SLs are important components of biological membranes. Besides their importance in cell integrity, they have drawn attention as bioactive molecules, playing a crucial role in cell function and fate (10). The SL metabolism consists of a network of various enzymes that catalyse the formation and degradation of the different SLs. Ceramide represents a central hub in this SL network. It can either be formed by *de novo* synthesis or by metabolism of complex SLs, e.g. sphingomyelin (11). Ceramidases hydrolyse ceramide to sphingosine, which can be further phosphorylated by sphingosine kinases, forming sphingosine-1-phosphate (S1P). So far, 5 different human ceramidases have been described according to their pH optimum: acid ceramidase (Ac, *ASAH1*), neutral ceramidase (Nc, *ASAH2*), and alkaline ceramidase 1-3 (Acer1-3, *ACER1-3*), each having a murine counterpart (12). Ac is localized in the lysosomes, where it preferentially hydrolyses ceramides with medium chain length (C12-C16) (13, 14). Mutations in the *ASAH1* gene can lead to two different rare diseases, the lysosomal storage disorder Farber disease (FD) and an epileptic disorder called spinal muscular atrophy with progressive myoclonic epilepsy (SMA-PME) (15). Ac and ceramide have also been implicated to play a role in several infectious diseases. While the pharmacological inhibition of Ac by ceranib-2 decreases measles virus replication *in vitro* (16), genetic overexpression of Ac rescues cystic fibrosis mice from pulmonary *Pseudomonas aeruginosa* infections *in vivo* (17). Lang et al. recently showed that Ac expression in macrophages limits the propagation of herpes simplex virus type 1 and protects mice from severe course of infection (18). However, the role of host cell-intrinsic Ac activity and ceramide during *Plasmodium* infection remains elusive.

Here, using the nonlethal rodent *Plasmodium* strain *P. yoelii* 17XNL as murine malaria model, we evaluated how Ac ablation and pharmacological inhibition modulate the course of infection in mice. We demonstrate that increased ceramide levels in Ac-deficient mice or treatment with the Ac inhibitor carmofur significantly reduce parasitemia in *P. yoelii*-infected mice. Mechanistically, we identified reduced generation of reticulocytes, the target cells of *P. yoelii*, upon deletion or inhibition of Ac.

## Material and Methods

### Mice

Mice were bred and housed under specific pathogen-free conditions at the animal facility of the University Hospital Essen. All experiments were performed in accordance with the guidelines of the German Animal Protection Law and approved by the State Agency for Nature, Environment, and Consumer Protection (LANUV), North Rhine-Westphalia, Germany (Az 84-02.04.2015.A474, Az 81-02.04.2018.A302). Female C57BL/6 mice were purchased from Envigo Laboratories (Envigo CRS GmbH). Ac^flox/flox^/creER mice (Asah1 tm1Jhkh/Gt(Rosa)26Sortm9(cre/ESR1)Arte) express a mutated form of the estrogen receptor (ER) which is fused to the cre recombinase (Taconic Biosciences Inc.). Two loxp-sites are flanking exon1 of the *Asah1* gene. Upon tamoxifen administration, the cre recombinase is translocated into nucleus leading to the deletion of exon1 and therefore loss of function. AC^flox/flox^/CD4cre mice (Asah1tm1Jhkh/Tg(CD4-cre)1Cwi) were generated by breeding AC^flox/flox^ mice with CD4cre mice (Tg(CD4-cre)1Cwi). The cre recombinase is expressed under the control of CD4 regulatory elements. AC^flox/flox^/LysMcre mice (Asah1tm1Jhkh/Lyz2tm1(Cre)Ifo) were obtained by crossing AC^flox/flox^ mice with LysMcre mice (Lyz2tm1(Cre)Ifo) that express cre recombinase under the control of the lysozyme M promoter (LysM).

### Tamoxifen administration

Tamoxifen (Sigma Aldrich) was dissolved in corn oil and administered to Ac^flox/flox^/creER^tg^ mice (Ac KO) and Ac^flox/flox^/creER^wt^ littermates (Ac WT) by intraperitoneal (i.p.) injection (4 mg in 100 µl) 8, 6, and 4 days prior to analysis or infection.

### Carmofur treatment

C57BL/6 mice were either treated with 750 µg carmofur (Abcam) dissolved in 100 µl corn oil or with corn oil only (vehicle) by daily i.p. injection starting 1 day prior to infection. For therapeutic use of carmofur, C57BL/6 mice were treated with carmofur or vehicle 5 days post infection (p.i.).

### *Plasmodium yoelii* infection

Cryopreserved *P. yoelii* 17XNL-infected red blood cells (iRBCs) were passaged once through C57BL/6 mice, before using them for infection of experimental mice. Experimental mice were infected with 1×10^5^ *P. yoelii*-parasitized red blood cells by intravenous (i.v.) injection on day 0. The parasitemia of infected mice was examined by microscopy of giemsa-stained blood films. RBC counts were analysed using the automated hematology analyzer KX-21N (Sysmex).

### Cell isolation and flow cytometry

Splenocytes were isolated by rinsing spleens with erythrocyte lysis buffer and washing with PBS supplemented with 2% FCS and 2 mM EDTA. Single cell suspensions of bone marrow cells were generated by flushing bones (tibia) with PBS. Blood was either taken by puncture of the tail vein during the experiment or by cardiac puncture at day of sacrifice, and immediately mixed with heparin to avoid coagulation. Magnetic activated cell sorting (MACS) was used to separate and enrich different cell populations from spleen, peritoneal lavage, or blood. Cells of interest were isolated using the CD4^+^/CD8a^+^ T cell isolation kit, the Neutrophil isolation kit, CD11c MicroBeads UltraPure, and F4/80 MicroBeads UltraPure (all Miltenyi Biotec) according to the manufacturer’s instructions and separated automatically with the AutoMACS from Miltenyi Biotec. CD4^+^ and CD8^+^ T cells were additionally sorted by fluorescence activated cell sorting (FACS) using an ARIA II cell sorter (BD Bioscience). For flow cytometry analyses, cells were stained with anti-CD4, anti-CD8, anti-CD49d (BD Bioscience), anti-Foxp3, anti-Ki67, anti-CD71 (eBioscience), anti-PD1, anti-CD11a, anti-CD45, and anti-Ter119 (Biolegend) conjugated to fluorescein isothiocyanate (FITC), pacific blue (PB), phycoerythrin (PE), allophycocyanin (APC), AlexaFlour647 (AF647), or PE-cyanin 7 (PE-Cy7). The fixable viability dye eFluor780 (eBioscience) was used to identify dead cells. Intracellular staining for Foxp3 and Ki67 was performed using the Fixation/Permeabilization kit (eBioscience) according to the manufacturer’s protocol. Cells were analysed by flow cytometry with a LSR II instrument (BD Bioscience) using DIVA software.

### Ceramide and sphingomyelin quantification by HPLC-MS/MS

Spleen tissue homogenates or cell suspensions were subjected to lipid extraction using 1.5 mL methanol/chloroform (2:1, v:v) as described before (19). The extraction solvent contained C17 ceramide (C17 Cer) and d31-C16 sphingomyelin (d31-C16 SM) (both Avanti Polar Lipids) as internal standards. Chromatographic separations were achieved on a 1290 Infinity II HPLC (Agilent Technologies) equipped with a Poroshell 120 EC-C8 column (3.0 × 150 mm, 2.7 µm; Agilent Technologies). MS/MS analyses were carried out using a 6495 triple-quadrupole mass spectrometer (Agilent Technologies) operating in the positive electrospray ionization mode (ESI+) (20). The following mass transitions were recorded (qualifier product ions in parentheses): <u>ceramides:</u> *m/z* 520.5 → 264.3 (282.3) for C16 Cer, *m/z* 534.5 → 264.3 (282.3) for C17 Cer, *m/z* 548.5 → 264.3 (282.3) for C18 Cer, *m/z* 576.6 → 264.3 (282.3) for C20 Cer, *m/z* 604.6 → 264.3 (282.3) for C22 Cer, *m/z* 630.6 → 264.3 (282.3) for C24:1 Cer, and *m/z* 632.6 → 264.3 (282.3) for C24 Cer; <u>sphingomyelins:</u> *m/z* 703.6 → 184.1 (86.1) for C16 SM, *m/z* 731.6 → 184.1 (86.1) for C18 SM, *m/z* 734.6→ 184.1 (86.1) for d31-C16 SM, *m/z* 759.6 → 184.1 (86.1) for C20 SM, *m/z* 787.7 → 184.1 (86.1) for C22 SM, *m/z* 813.7 → 184.1 (86.1) for C24:1 SM, and *m/z* 815.7 → 184.1 (86.1) for C24 SM. Peak areas of Cer and SM subspecies, as determined with MassHunter software (Agilent Technologies), were normalized to those of the internal standards (C17 Cer or d31-C16 SM) followed by external calibration in the range of 1 fmol to 50 pmol on column. Determined Cer and SM amounts were normalized either to cell count or to the actual protein content (as determined by Bradford assay) of the tissue homogenate used for extraction.

### RNA isolation, cDNA synthesis, and qRT-PCR

RNA was either isolated using the RNeasy Fibrous Tissue Kit (Qiagen) for spleen, thymus, and liver tissue, or using the RNeasy Mini Kit (Qiagen) for bone marrow cells, T cells, macrophages, dendritic cells, and neutrophils according to the manufacturer’s instructions. To synthesize cDNA, 0.1-1 µg of RNA was reversed transcribed using M-MLV Reverse Transcriptase (Promega) with dNTPs, Oligo-dT mixed with Random Hexamer primers (Thermo Fisher Scientific). For quantitative real-time PCR Fast SYBR Green Master Mix (Thermo Fisher Scientific) was used on a 7500 Fast Real-Time PCR System (Thermo Fisher Scientific). Each sample was measured as technical duplicate. The expression levels of target genes were normalized against *ribosomal protein S9* (*RPS9*). Following primer sequences (5’-3’) were used: *Asah1* (TTC TCA CCT GGG TCC TAG CC, TAT GGT GTG CCA CGG AAC TG), *Asah2* (AGA GAG AGC AAG GTA TTC TTC, ACT ATT CAC AAA GTG GTT GC), *RPS9* (CTG GAC GAG GGC AAG ATG AAG C, TGA CGT TGG CGG ATG AGC ACA).

### Statistical analyses

Statistical analyses were performed using GraphPad Prism 7 software. To test for Gaussian distribution, D’Agostino-Pearson omnibus and Shapiro-Wilk normality tests were used. If data passed normality testing unpaired Student’s t-test was performed, otherwise Mann-Whitney U-test was used. Differences between two or more groups with different factors were calculated using two-way ANOVA followed by Sidak’s post-test. *P*-values of 0.05 or less were considered indicative of statistical significance (\**p* < 0.05, \*\**p* < 0.01, \*\*\**p* < 0.001, \*\*\*\**p* < 0.0001).

## Results

### Ac-deficient mice accumulate ceramide

Since the genetic ablation of Ac is embryonically lethal in mice, we made use of inducible Ac^flox/flox^/creER^tg^ (Ac KO) mice, which received tamoxifen 8, 6, and 4 days prior to analysis or *P. yoelii* infection (Fig. 1A) (21, 22). Tamoxifen-treated Ac^flox/flox^/creER^wt^ (Ac WT) mice littermates served as control. As depicted in Fig. 1B, Ac mRNA expression was ubiquitously abolished in all analysed tissues in Ac KO mice. Consequently, ceramide concentrations determined by mass spectrometry were significantly increased in spleens from Ac KO mice compared to Ac WT mice (Fig. 1C).

**Fig. 1:**
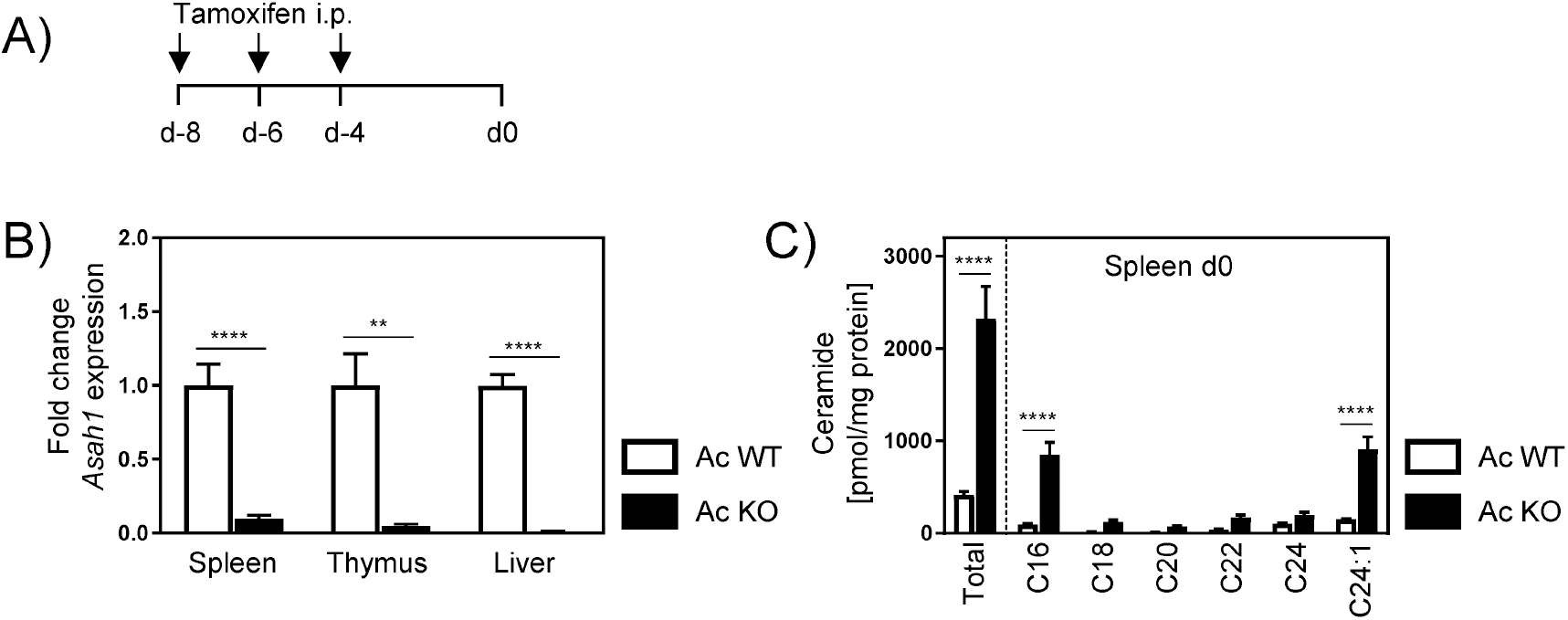
Ceramide accumulation in Ac KO mice. (A) Induction of Ac deficiency in Ac^flox/flox^/creER mice (Ac KO) was achieved by intraperitoneal injection (i.p.) of 4 mg tamoxifen on day −8, −6, and −4. Tamoxifen-treated wild type littermates (Ac WT) were used as control. (B) Ac KO validation was performed by analysing *Asah1* mRNA expression via RT-qPCR in spleen, thymus, and liver of Ac KO mice and Ac WT littermates as control at day 0 (n=5-6). (C) Ceramide levels in spleen were determined by HPLC-MS/MS (n=3-4). Data are presented as mean ± SEM. Statistical analyses were performed using Mann-Whitney U-test and two-way ANOVA, followed by Sidak’s post-test (\*\**p*<0.01, \*\*\*\**p*<0.0001).

### Ac deficiency results in reduced parasitemia and decreased T cell responses during early stage of *P. yoelii* infection

To study the role of Ac during malaria, we infected Ac KO mice and Ac WT littermates with *P. yoelii*. Interestingly, Ac KO mice showed significantly reduced parasitemia at day 3 and 7 p.i. (Fig. 2A). However, these differences were no longer detectable 10 and 14 days p.i.. As splenomegaly is a common clinical signature of *Plasmodium* infections, we determined the spleen weight of infected Ac WT and Ac KO mice. Well in line with the observed effect on parasitemia, spleen weights of Ac KO mice were tremendously decreased compared to those of Ac WT mice 7 days p.i., but no longer differed at day 14 p.i. (Fig. 2B).

**Fig. 2:**
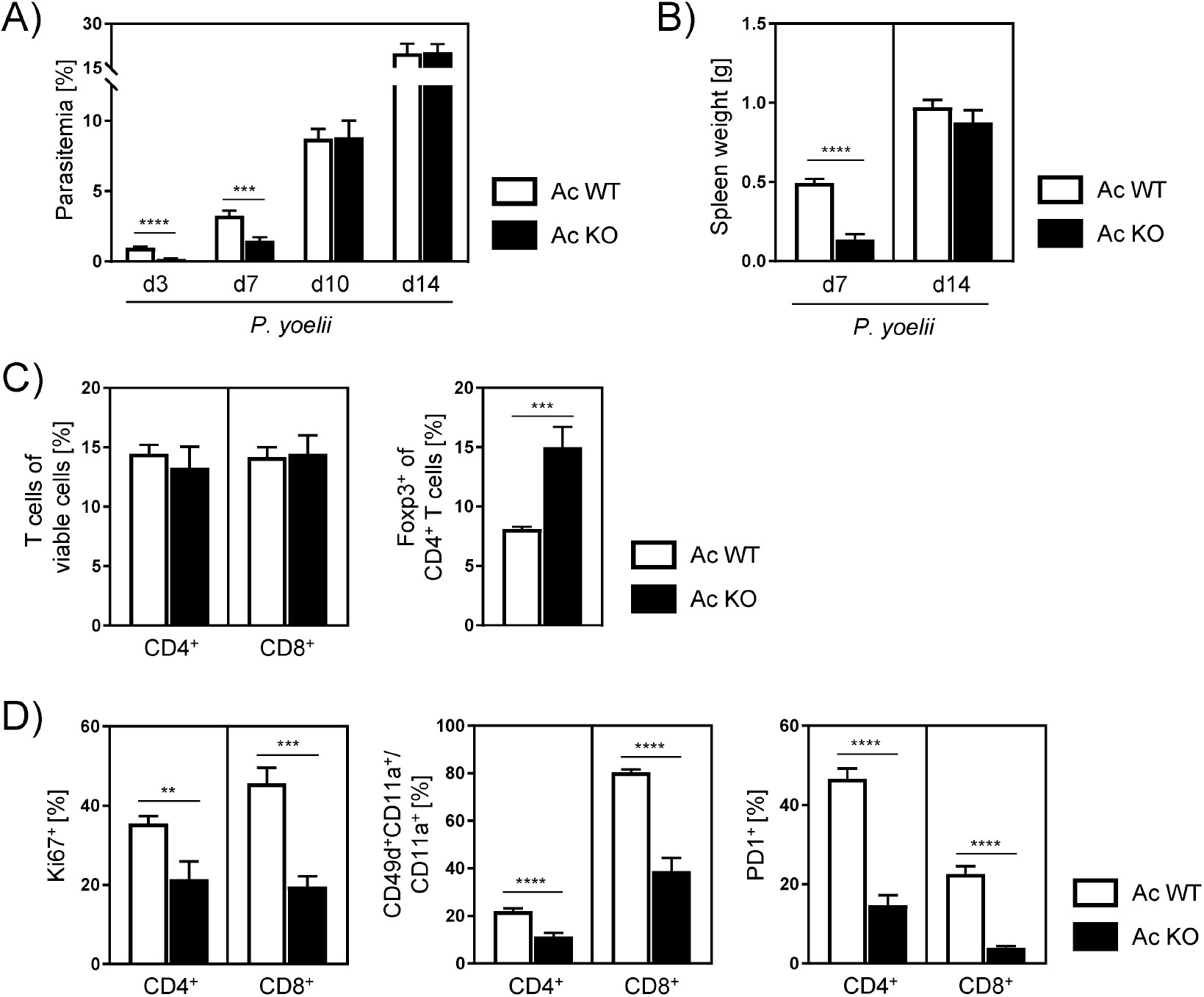
*P. yoelii*-infected Ac KO mice show decreased parasitemia with less T cell activation in the early phase of infection. (A) Parasitemia of *P. yoelii*-infected Ac WT and Ac KO mice was determined at indicated time points by microscopy of giemsa-stained blood films (n=9-18). (B) Spleen weight on day 7 and 14 post infection (p.i.) (n=7-9). (C) Frequencies of viable CD4^+^, CD8^+^, and regulatory T cells (Foxp3^+^ of CD4^+^) were analysed by flow cytometry 7 days p.i. (n=7-9). (D) Percentages of Ki67-, CD49dCD11a/CD11a-, and PD1-expressing CD4^+^ and CD8^+^ T cells were determined by flow cytometry 7 days p.i. (n=7-9). Data from three to five independent experiments are presented as mean ± SEM. Statistical analyses were performed using unpaired Student’s t-test or Mann-Whitney U-test (\*\**p*<0.01, \*\*\**p*<0.001, \*\*\*\**p*<0.0001).

Next, we analysed the phenotype of T cells from spleen of infected mice 7 days p.i. by flow cytometry. T cells, especially CD4^+^ T cells, have been described to play an important role in limiting blood-stage malaria (8). While frequencies of CD4^+^ and CD8^+^ T cells did not differ between *P. yoelii*-infected Ac WT and Ac KO mice, we found significantly increased frequencies of CD4^+^Foxp3-expressing regulatory T cells (Tregs) in spleen of Ac KO mice compared to Ac WT littermates (Fig. 2C). Interestingly, CD4^+^ and CD8^+^ T cells from Ac KO mice showed decreased expression of marker molecules associated with T cell activation, namely Ki67, CD49d, CD11a, and PD1 (Fig. 2D). These results show that Ac deficiency leads to reduced parasitemia in the early phase of *P. yoelii* infection accompanied by diminished T cell responses.

### Cell-specific Ac deletion in T cells and myeloid cells does not affect the course of *P. yoelii* infection

In order to further evaluate the effect of Ac on immune responses during *P. yoelii* infection, we made use of cell type-specific Ac-deficient mice. Since T cells from infected global Ac KO mice were less activated, we next analysed Ac^flox/flox^/CD4cre mice that specifically lack Ac expression in CD4^+^ and CD8^+^ T cells (Fig. 3A). However, T cell-specific Ac ablation had no impact on the course of *P. yoelii* infection, as Ac CD4cre KO (Ac^flox/flox^/CD4cre^tg^) and Ac CD4cre WT (Ac^flox/flox^/CD4cre^wt^) mice showed similar spleen weights and parasitemia 7 and 14 days p.i. (Fig. 3B). We were wondering whether an early enhanced innate immune response as first line of defence contribute to an improved parasite clearance in global Ac KO mice and therefore to less T cell activation in the initial phase of infection. Thus, we used Ac^flox/flox^/LysMcre mice, which exhibit decreased Ac expression in myeloid cells, e.g. macrophages, dendritic cells, and neutrophils (Fig. 3C). However, upon *P. yoelii* infection, Ac LysMcre KO (Ac^flox/flox^/LysMcre^tg^) mice and Ac LysMcre WT (Ac^flox/flox^/LysMcre^wt^) littermates did not significantly differ regarding parasitemia and spleen weight (Fig. 3D).

**Fig. 3:**
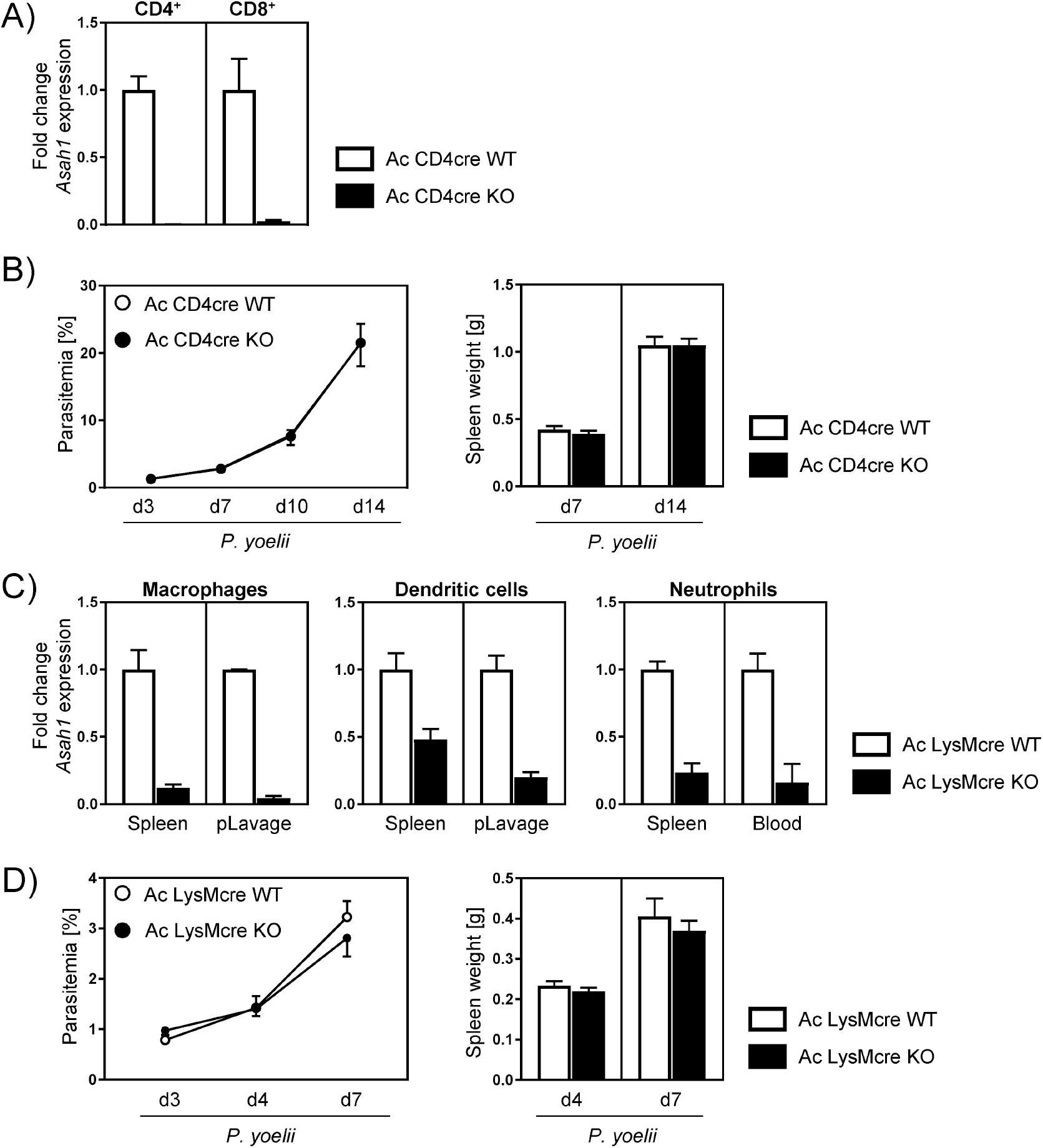
T cell-specific and myeloid-specific Ac deletion has no impact on the course of *P. yoelii* infection. (A) The knockout of Ac in T cells was confirmed by analysing *Asah1* mRNA expression of sorted splenic CD4^+^ and CD8^+^ T cells from naïve Ac^flox/flox^/CD4cre^tg^ (Ac CD4cre KO) mice and Ac^flox/flox^/CD4cre^wt^ littermates (Ac CD4cre WT) as controls via RT-qPCR (n=2-4). (B) Parasitemia (left panel) and spleen weight (right panel) of *P. yoelii*-infected Ac CD4cre KO mice and Ac CD4cre WT littermates was determined at indicated time points (n=7-10). (C) The knockout of Ac in myeloid cells was confirmed by analysing *Asah1* mRNA expression of macrophages, dendritic cells, and neutrophils isolated from spleen, peritoneal lavage (pLavage), and blood of naïve Ac^flox/flox^/LysMcre^tg^ (Ac LysMcre KO) mice and Ac^flox/flox^/LysMcre^wt^ littermates (Ac LysMcre WT) as controls via RT-qPCR (n=2-6). (D) Parasitemia (left panel) and spleen weight (right panel) of *P. yoelii*-infected Ac LysMcre KO and Ac LysMcre WT mice was determined at indicated time points (n=9). Data from two independent experiments each are presented as mean ± SEM.

### Ac KO mice show an impaired erythropoiesis

Since global Ac KO mice showed decreased parasitemia, although cell-specific Ac activity did not alter innate and T cell immune responses during *P. yoelii* infection *per se*, we analysed whether Ac deletion instead has an effect on the host cells of the parasites. Depending on the species, *Plasmodium* parasites prefer immature or mature red blood cells for asexual propagation (23). *P. yoelii* parasites primarily infect reticulocytes, which are defined as Ter119^+^CD71^+^ (24). Therefore, we analysed frequencies of reticulocytes in blood of Ac WT and Ac KO mice by flow cytometry. Strikingly, reticulocytes were dramatically reduced in blood of non-infected Ac KO mice (Fig. 4A, d0). While they were almost absent on day 0 and 3 p.i., frequencies rapidly increase in Ac KO mice7 days p.i. resulting in equal percentages of reticulocytes in Ac WT and Ac KO mice 10 days p.i. (Fig. 4B).

**Fig. 4:**
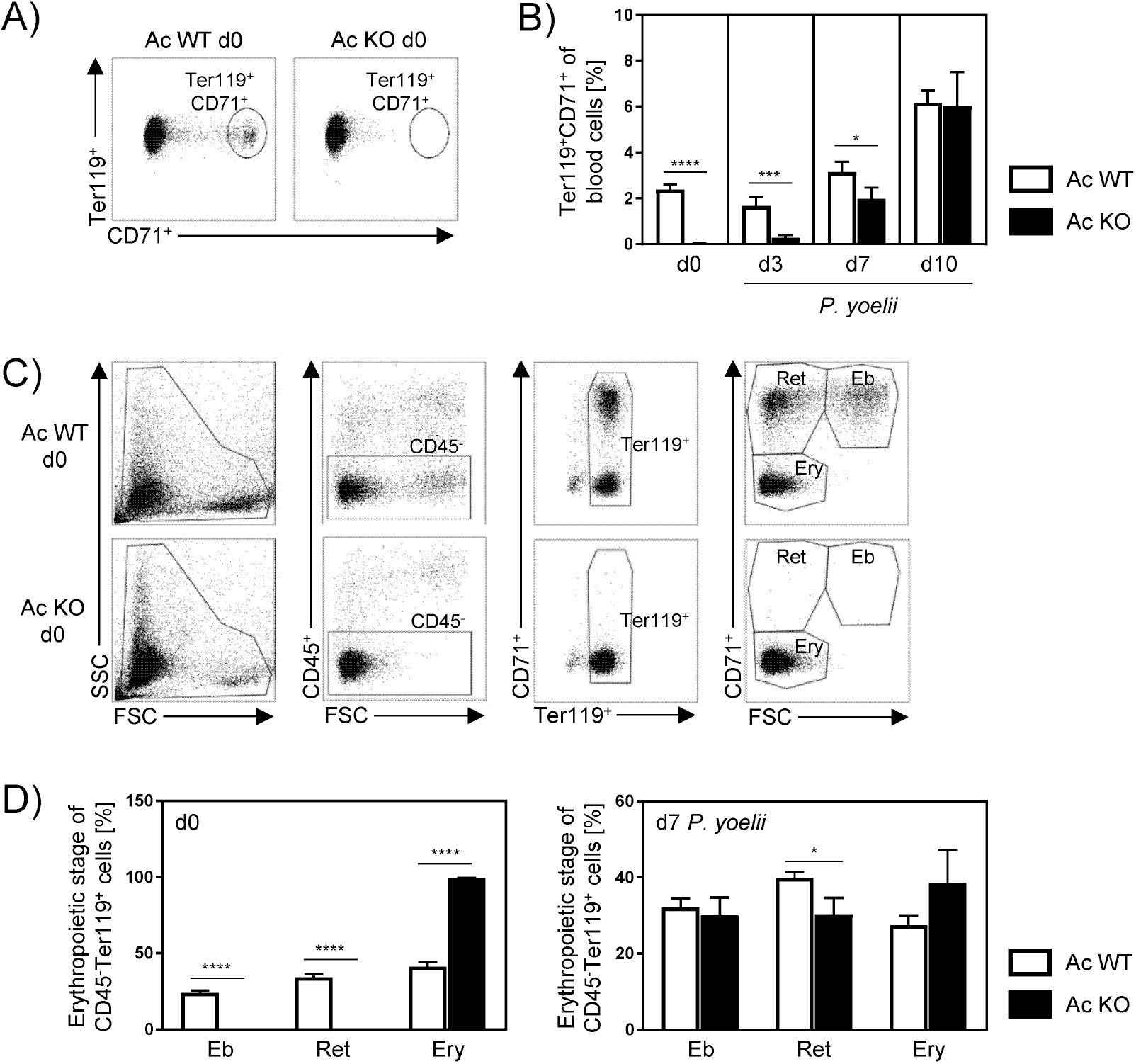
Ac KO affects the erythropoiesis. (A) Representative flow cytometry dot plot of Ter119^+^CD71^+^ reticulocytes in blood of Ac WT and Ac KO mice on day 0. (B) Frequencies of reticulocytes in blood of Ac WT and Ac KO mice were determined by flow cytometry on day 0 and after *P. yoelii* infection on day 3, 7, and 10 (n=8-15). (C) Representative flow cytometry gating strategy of erythropoietic developmental stages in bone marrow of Ac WT and Ac KO mice on day 0. Erythropoietic cells were defined as CD45^-^ and Ter119^+^. Depending on size (forward scatter, FSC) and CD71 expression, cells were defined as erythroblasts (Eb, CD71^+^FSC^high^), reticulocytes (Ret, CD71^+^FSC^low^), and mature erythrocytes (Ery, CD71^-^FSC^low^). (D) Frequencies of different erythropoietic stages in bone marrow of non-infected Ac WT and Ac KO mice on day 0 (left panel) and after *P. yoelii* infection on day 7 (right panel) were analysed by flow cytometry (n=6-11). Results from two to four independent experiments are presented as mean ± SEM. Statistical analyses were performed using unpaired Student’s t-test or Mann-Whitney U-test (\**p*<0.05, \*\*\**p*<0.001, \*\*\*\**p*<0.0001).

To gain further insights into the impact of Ac expression on erythropoietic cells, we analysed the development of RBCs within the bone marrow of Ac WT and Ac KO mice in more detail. Depending on size and CD71 expression, erythropoietic cells (CD45^-^ Ter119^+^) were identified as erythroblasts, reticulocytes, and mature erythrocytes (Fig. 4C). As depicted in Fig. 4D, erythroblasts and reticulocytes were hardly detectable in bone marrow of non-infected Ac KO at day 0 (left panel). Consistent with raising reticulocyte frequencies in the blood (Fig. 4B), percentages of erythroblasts and reticulocytes were increasing 7 days p.i. in bone marrow of *P. yoelii*-infected Ac KO mice, but they were still reduced when compared to infected Ac WT mice (Fig. 4D, right panel). Similar results were obtained in non-infected Ac KO mice (Fig. 4 – figure supplement 1), providing evidence that this phenotype is independent of the infection. Together, these results indicate an impaired erythropoiesis in Ac KO mice, leading to reduced frequencies of reticulocytes the main target cells of *P. yoelii* parasites.

### Ablation of Ac also affects sphingomyelin levels

Our data show that Ac deficiency alters erythropoiesis, resulting in decreased frequencies of reticulocytes in peripheral blood and bone marrow of Ac KO mice. Nonetheless, during *P. yoelii* infection of Ac KO mice reticulocytes start to repopulate, which is also reflected in rapidly rising parasitemia 10 and 14 days p.i. (Fig. 2A). To confirm that the induced Ac KO is persistent at later time points, we analysed Ac expression in the bone marrow of non-infected mice at day 0 and 7 and 10 days after *P. yoelii* infection. As validated by RT-qPCR, Ac expression was still absent in Ac KO mice 7 and 10 days p.i. (Fig. 5A).

**Fig. 5:**
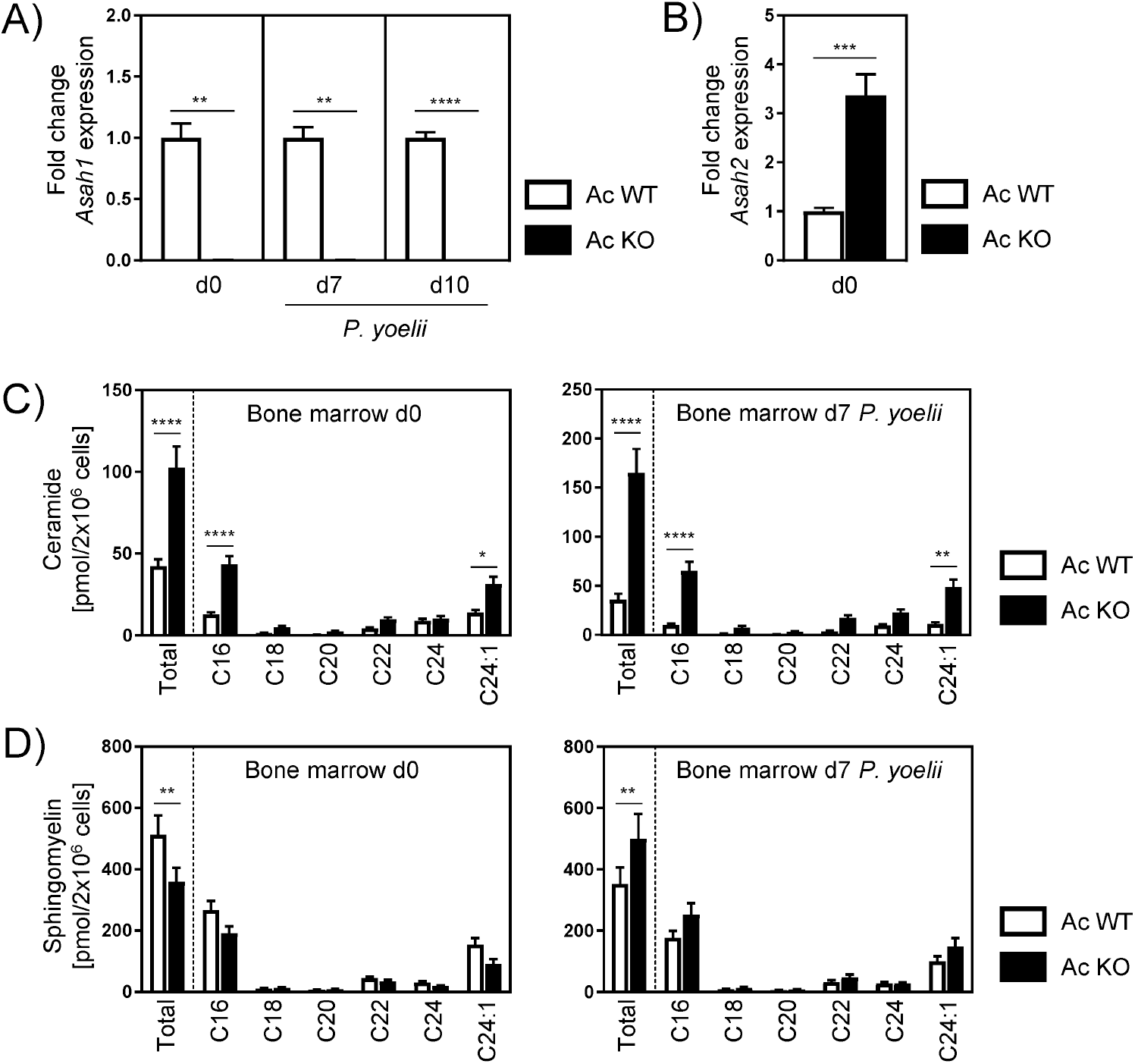
Alterations of sphingolipid metabolism in bone marrow of Ac KO mice. (A) Ac KO validation in bone marrow of non-infected (d0) and *P. yoelii*-infected (d7 and d10) Ac KO mice and Ac WT littermates as control by RT-qPCR (n=5-9). (B) Fold change of neutral ceramidase (*Asah2*) expression in bone marrow at day 0 (n=6-7). (C) Ceramide and (D) sphingomyelin levels in bone marrow of non-infected Ac WT and Ac KO on day 0 (left panel), and 7 days p.i. (right panel) were determined by HPLC-MS/MS (n=3-6). Data are presented as mean ± SEM. Statistical analyses were performed using unpaired Student’s t-test and two-way ANOVA, followed by Sidak’s post-test (\**p*<0.05, \*\**p*<0.01, \*\*\**p*<0.001, \*\*\*\**p*<0.0001).

In order to investigate whether other ceramidases compensate for Ac activity, we analysed their mRNA expression. We found the neutral ceramidase (Nc) to be upregulated after induction of the Ac KO (Fig. 5B). However, analysis of ceramide levels in bone marrow of Ac WT and Ac KO mice revealed significantly higher ceramide concentrations in bone marrow of Ac KO mice compared to Ac WT littermates 7 days p.i. (Fig. 5C, right panel), similar to those measured at day 0 (Fig. 5C left panel). This suggests that elevated Nc expression in Ac KO mice is not sufficient to completely compensate for Ac deficiency with regard to the ceramide content.

Next, we determined sphingomyelin levels in bone marrow of Ac WT and Ac KO mice. Sphingomyelin is generated from ceramide by the activity of sphingomyelin synthases and has been shown to increase the number of reticulocytes in mice (25). While sphingomyelin levels were decreased in bone marrow of Ac KO mice compared to Ac WT mice at day 0 (Fig. 5D, left panel), they were significantly enhanced at day 7 in presence (Fig. 5D, right panel) and absence (Fig. 5 – figure supplement 1) of the infection, correlating with increasing reticulocyte frequencies observed before. These results indicate an altered SL metabolism in bone marrow of Ac KO mice that does not only affect ceramide levels but also leads to elevated sphingomyelin concentrations 7 days p.i., which might account for the repopulation of reticulocytes.

### Carmofur treatment leads to alleviated parasitemia and reduced reticulocyte frequencies during *P. yoelii* infection

Since the genetic ablation of Ac strongly affected erythropoiesis and, as a consequence the course of *P. yoelii* infection, we investigated whether pharmacological inhibition of Ac has the same effect. Studies by Realini et al. showed that the antineoplastic drug carmofur is a potent inhibitor of Ac activity in mice (26). Therefore, C57BL/6 mice were treated with carmofur or vehicle and infected with *P. yoelii* (Fig. 6A). Strikingly, carmofur-treated mice showed a strong reduction in parasitemia (Fig. 6B). Moreover, spleen weights of carmofur-treated mice were significantly reduced at day 7 p.i. (Fig. 6C).

**Fig. 6:**
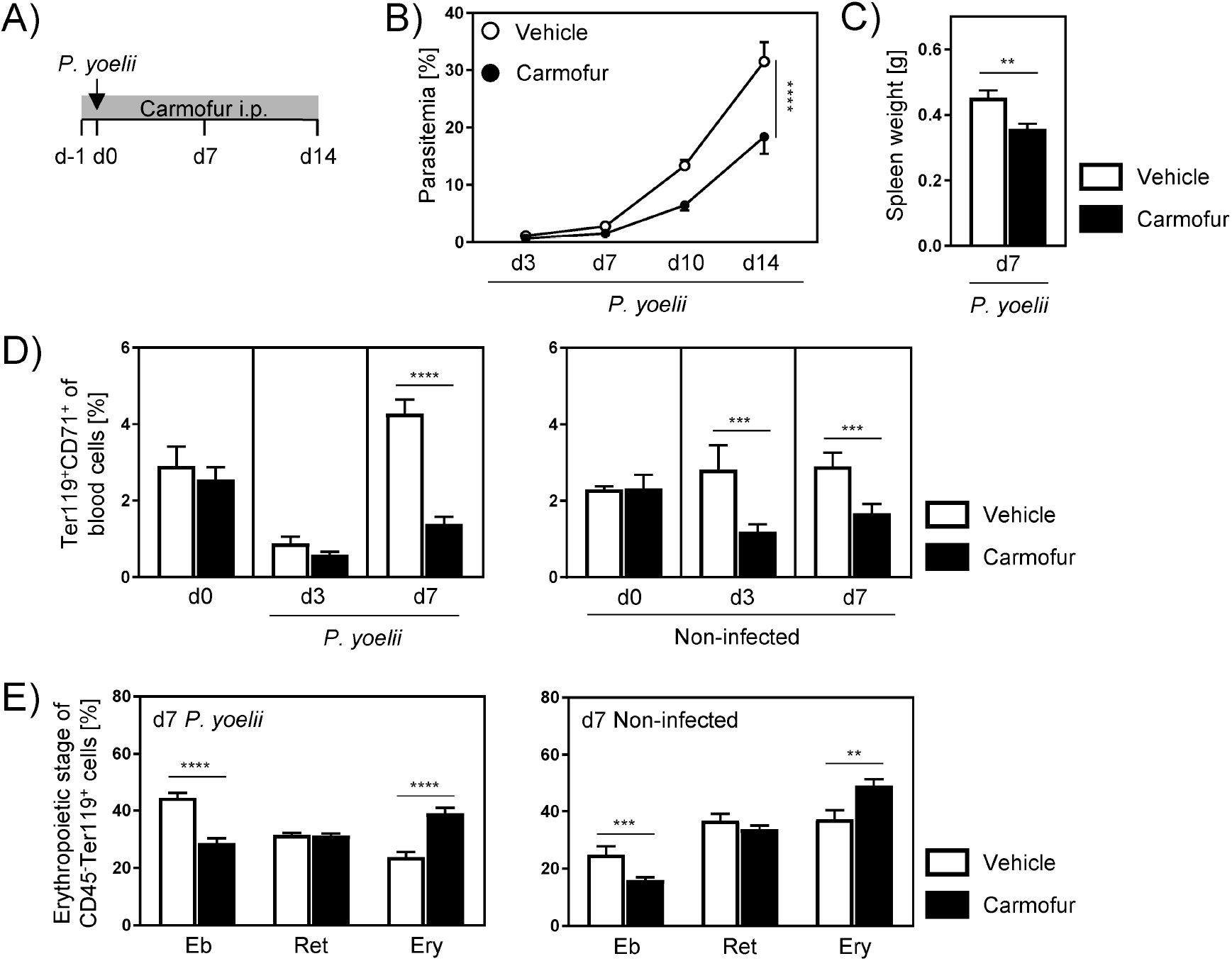
Carmofur treatment leads to decreased parasitemia with lower reticulocyte frequencies. (A) For pharmacological inhibition of Ac, C57BL/6 mice were either treated with 750 µg carmofur or with vehicle as control by daily intraperitoneal (i.p.) injection starting one day prior to *P. yoelii* infection. (B) Parasitemia of infected carmofur- and vehicle-treated mice was determined at indicated time points by microscopy of giemsa-stained blood films (n=10-11). (C) Spleen weight of *P. yoelii*-infected carmofur- and vehicle-treated mice 7 days p.i. (n=9). (D) Frequencies of Ter119^+^CD71^+^ reticulocytes in blood of *P. yoelii*-infected (left panel) and non-infected (right panel) carmofur- and vehicle-treated mice were analysed by flow cytometry on day 0, 3, and 7 (n=10-14). (E) Percentages of erythroblasts (Eb, CD71^+^FSC^high^), reticulocytes (Ret, CD71^+^FSC^low^), and mature erythrocytes (Ery, CD71^-^FSC^low^) in bone marrow of *P. yoelii*-infected (left panel) and non-infected (right panel) carmofur- and vehicle-treated mice on day 7 (n=9). Results from two to three independent experiments are presented as mean ± SEM. Statistical analyses were performed using two-way ANOVA, followed by Sidak’s post-test, unpaired Student’s t-test or Mann-Whitney U-test (\*\**p*<0.01, \*\*\**p*<0.001, \*\*\*\**p*<0.0001).

In order to analyse whether the milder course of infection in carmofur-treated mice is also associated with altered reticulocyte frequencies, we analysed blood of carmofur-and vehicle-treated mice at day 0 as well as 3 and 7 days after *P. yoelii* infection. While there is a tendency of decreased reticulocytes on day 0 and 3, carmofur-treated mice showed significantly reduced reticulocyte frequencies compared to vehicle-treated mice at day 7 p.i. (Fig. 6D, left panel). Furthermore, carmofur treatment led to an altered erythropoiesis in bone marrow with lower relative numbers of erythroblasts compared to vehicle control 7 days p.i. (Fig. 6E, left panel). Similar results were obtained in non-infected carmofur-and vehicle-treated mice (Fig. 6D and 6E, right panels), indicating that modulation in erythropoiesis occurs independent from the infection.

### Carmofur as efficient therapy against *P. yoelii* infection

Since therapy in clinics is usually started after the infection has already manifested, we next analysed the therapeutic potential of carmofur treatment. Therefore, we infected C57BL/6 mice with *P. yoelii* and started with carmofur treatment at day 5 p.i., when the infection was established (Fig. 7A). As depicted in Fig. 7B, mice showed significantly reduced reticulocyte frequencies after carmofur therapy 7, 10, and 14 days p.i.. This was accompanied by a strongly alleviated parasitemia (Fig. 7C). Moreover, spleen weights of carmofur-treated mice were decreased compared to vehicle control (Fig. 7D). Although carmofur therapy seem to reduce the production of reticulocytes, carmofur-treated mice showed similar or even slightly higher RBC counts than vehicle-treated mice during infection, indicating they did not suffer from a more severe anemia (Fig. 7E).

**Fig. 7:**
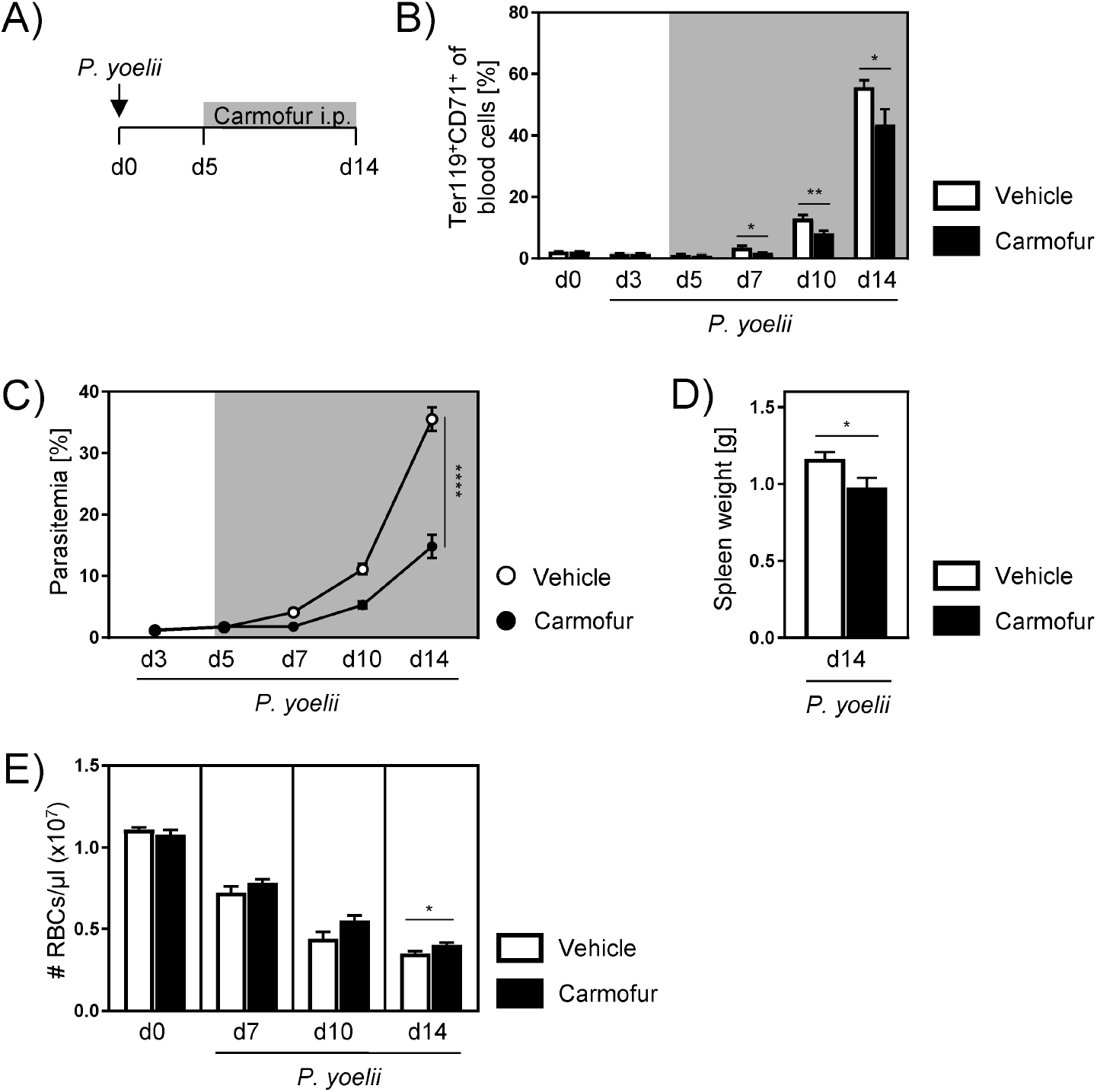
Carmofur application as therapeutic treatment of *P. yoelii* infection. (A) As therapeutic approach, C57BL/6 mice were infected with *P. yoelii* on day 0 and treated with 750 µg carmofur or vehicle only from day 5 p.i. onwards. (B) Percentages of Ter119^+^CD71^+^ reticulocytes in blood were analysed by flow cytometry at indicated time points (n=5-10). (C) Parasitemia of infected carmofur- and vehicle-treated mice was determined 3, 5, 7, 10, and 14 days p.i. by microscopy of giemsa-stained blood films (n=10). (D) Spleen weight of *P. yoelii*-infected carmofur- and vehicle-treated mice 14 days p.i. (n=10). (E) Red blood cell (RBC) count per microliter blood of infected carmofur- and vehicle-treated mice was determined with an automated hematology analyzer (n=5-10). Results from two independent experiments are summarized as mean ± SEM. Statistical analyses were performed using unpaired Student’s t-test, Mann-Whitney U-test, or two-way ANOVA, followed by Sidak’s post-test, or (\**p*<0.05, \*\**p*<0.01, \*\*\*\**p*<0.0001).

In summary, our results provide evidence that deficiency or inhibition of Ac activity alters erythropoiesis, resulting in decreased reticulocyte frequencies associated with lower parasitemia during *P. yoelii* infection.

## Discussion

By hydrolysing ceramide into sphingosine, Ac is a key regulator of the SL metabolism. We and others showed that ceramide signalling is involved in the regulation of cell differentiation, activation, apoptosis, and proliferation (9, 27, 28). However, the impact of cell-intrinsic Ac activity and ceramide on the course of *Plasmodium* infection remains unclear. Here, we made use of Ac-deficient mice with ubiquitously increased ceramide levels to study the impact of endogenous ceramide on *P. yoelii* infection.

An adequate immune response is necessary to fight malaria-causing *Plasmodium* infections. It is well known, that T cells play an important role during the blood-stage of malaria (8). Our data show decreased expression of molecules associated with T cell activation in Ac KO mice in the early infection phase (Fig. 2D). However, this observation is rather due to a low parasite load, than due to reduced T cell activation caused by Ac-deficiency *per se*. As T cell-specific Ac ablation in Ac CD4cre KO mice did not affect the course of *P. yoelii* infection (Fig. 3B), endogenous ceramide levels do not seem to have a significant impact on T cell activation in this model.

In addition to its characteristics mentioned before, ceramide has also been proposed to modulate erythropoiesis. While long chain ceramides (C22, C24:0, and C24:1) are described to induce erythroid differentiation in bone marrow cells of rats (29), the treatment with C2 ceramide or bacterial ceramide-generating sphingomyelinase resulted in the inhibition of erythropoiesis in human bone marrow cells (30). However, these studies were conducted with externally applied ceramide and exclusively *in vitro*. The role of cell-intrinsic ceramide on RBC development *in vivo* largely remained elusive. Our data indicate that loss of Ac activity and consequently increased ceramide levels dramatically affect erythrocyte maturation *in vivo*. Bone marrow of Ac KO mice contained almost no erythroblasts and reticulocytes at day 0 (Fig. 4D, left panel). Well in line, studies by Orsini et al. showed reduced frequencies of orthochromatophilic erythroblasts and absence of reticulocytes in a culture of human hematopoietic stem progenitor cells from umbilical cord blood in response to C2 ceramide or bacterial sphingomyelinase treatment (31). They propose an inhibition of erythropoiesis independent of apoptosis, mediated by the induction of myelopoiesis via the TNFα/neutral sphingomyelinase/ceramide pathway. In contrast, Signoretto and colleagues observed increased annexin-V staining and therefore elevated eryptosis of human erythrocytes in response to ceramide accumulation induced by treatment with the ceramidase inhibitor ceranib-2 (32). Although we did not detect a shift from erythropoiesis towards myelopoiesis at day 0 (data not shown), RBCs from Ac KO mice exhibited significantly reduced annexin-V staining compared to Ac WT controls after 24 and 72 h of *in vitro* incubation (data not shown). Thus, our data suggest that lower reticulocyte frequencies are not due to increased apoptosis of RBCs. Nevertheless, erythroblasts and reticulocytes seem to repopulate bone marrow and blood of *P. yoelii*-infected Ac KO mice from day 7 p.i. on (Fig. 4B and 4D, right panel). Since ceramide levels continued to increase until day 7 p.i. (Fig. 5C), there was at least no sufficient compensation by other ceramidases, although mRNA expression of Nc was upregulated at day 0 (Fig. 5B). However, while sphingomyelin levels were significantly reduced in bone marrow of Ac KO mice compared to Ac WT littermates at day 0 (Fig. 5D, left panel), they were highly elevated at day 7 p.i. (Fig. 5D, right panel) when erythroblasts and reticulocytes reappear. Sphingomyelin is generated from ceramide by sphingomyelin synthases, which are important regulators of intracellular and plasma membrane ceramide and sphingomyelin levels (33). Consistent with our data, studies by Clayton et al. showed an induction of erythropoiesis by sphingomyelin in bone marrow of rats *in vitro* (29). Moreover, Scaro and colleagues observed an increase of circulating reticulocytes after injecting mice with purified sphingomyelin *in vivo* (25). Hence, one might speculate that the massive ceramide accumulation in Ac KO mice leads to an augmented SL turnover, resulting in elevated sphingomyelin concentrations in the bone marrow of Ac KO mice 7 days p.i. that might be responsible for resumption of erythropoiesis.

Several studies provide evidence that SLs and SL analogues are capable of regulating the development of *Plasmodium* parasites (34, 35). C6 ceramide has been shown to efficiently inhibit growth of *P. falciparum* parasites in a dose-dependent manner *in vitro* by reducing cytosolic glutathione levels and thereby inducing death of the parasites (36). This cytotoxic effect was abolished by addition of S1P, a downstream metabolite of ceramide and sphingosine, respectively. The importance of S1P for parasite survival has recently also been analysed by Sah et al. (37). They demonstrated that *P. falciparum* parasites strictly depend on the erythrocyte endogenous S1P pool for their intracellular development. Inhibition of the S1P-generating enzyme sphingosine kinase-1 resulted in decreased glycolysis, which is essential for parasite energy supply. Although we could not exclude an impact of erythrocytic glutathione or S1P levels, it is more likely that the lower parasite load in the early phase of infection is due to highly reduced reticulocyte frequencies in blood of Ac KO mice (Fig. 4B), which represent the target cells of *P. yoelii* parasites (24). Moreover, after reappearance of the reticulocytes, Ac KO an Ac WT mice exhibited similar parasitemia 10 and 14 days p.i. (Fig. 2A).

As our data indicate the involvement of the Ac/ceramide axis in regulating erythropoiesis, which affects the course of *Plasmodium* infection, we speculated that these findings might open the door to new potential therapeutic strategies against malaria. The limited availability of antimalarial drugs and emerging parasite resistances demand the development of new approaches. Therefore, we made use of carmofur, an antineoplastic drug that has recently been described to inhibit Ac activity *in vitro* and *in vivo* (26, 38). Carmofur-treated mice exhibited significantly reduced parasitemia and spleen weight compared to control mice during *P. yoelii* infection (Fig. 6B and 6C). Consistent with our observations in Ac KO mice, carmofur treatment led to decreased reticulocyte frequencies and an altered RBC development (Fig. 6D and 6E). Previous studies have shown the induction of reticulocytosis during the course of *P. yoelii* infection (24, 39). As we cannot exclude a direct cytotoxic effect of carmofur on the parasites, the question arises whether the lower parasite load in our model is due to the reduced reticulocyte count or, conversely, whether the lower parasitemia causes the lower reticulocytosis. Although it does not completely preclude the effect of carmofur on the parasites, non-infected carmofur-treated mice show similar changes in erythropoiesis that we observed in the infected animals (Fig. 6D and 6E, left panels).

In contrast to the murine model of blood-stage malaria, human *Plasmodium* infections often go along with low erythropoiesis and reticulocytosis, respectively (40, 41). On the one hand, especially for reticulocyte-restricted species as *P. vivax*, this leads to a limited number of target cells for the parasites. On the other hand, it results in anaemia that causes mortality and morbidity in patients. However, studies by Chang et al. and mathematical calculations by Cromer and colleagues revealed that these beneficial or detrimental effects depend on timing and the rate of reticulocyte production (42, 43). Although carmofur-treated mice show a lower reticulocyte production and therefore lower parasitemia, they had similar or even higher RBC counts compared to vehicle-treated controls, meaning they did not suffer from a more severe anaemia (Fig. 7E). Thus, carmofur treatment seem to beneficially influence the balance between parasite burden and RBC production. Finally, therapeutic carmofur administration to *P. yoelii*-infected mice also led to a decreased parasitemia and spleen weight (Fig. 7), demonstrating the efficacy of carmofur therapy against malaria, even after manifestation of infection.

Overall, our results indicate that cell-intrinsic Ac activity plays an important role during erythropoiesis and therefore during blood-stage infections. Hence, pharmacological inhibition might represent a novel therapeutic strategy to conquer malaria and other infections caused by reticulocyte-prone pathogens.

## Acknowledgement

We kindly thank Sina Luppus and Christina Liebig for excellent technical assistance. We further thank Daniel Herrmann for the help with HPLC-MS/MS sphingolipid quantification. This work was supported by the Deutsche Forschungsgemeinschaft (DFG - GRK2098 to AMW, JB, KSL, EG, WH, and DFG - GRK2581 to BK).

## Figure legends

**Fig. 4 – figure supplement 1:**
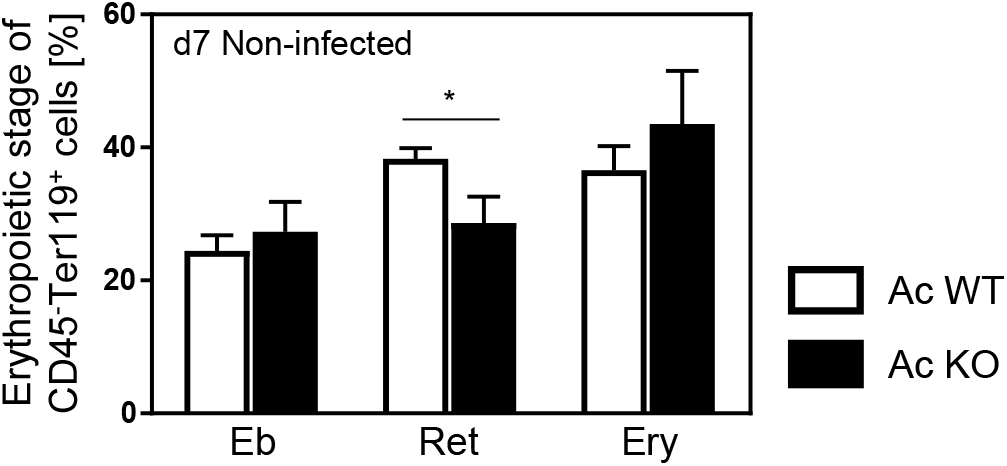
Erythropoiesis of non-infected Ac KO and Ac WT mice. Frequencies of different erythropoietic stages in bone marrow of non-infected Ac WT and Ac KO mice on day 7 were analysed by flow cytometry (n=12). Data from three independent experiments are presented as mean ± SEM. Statistical analyses were performed using unpaired Student’s t-test (\**p*<0.05).

**Fig. 5 – figure supplement 1:**
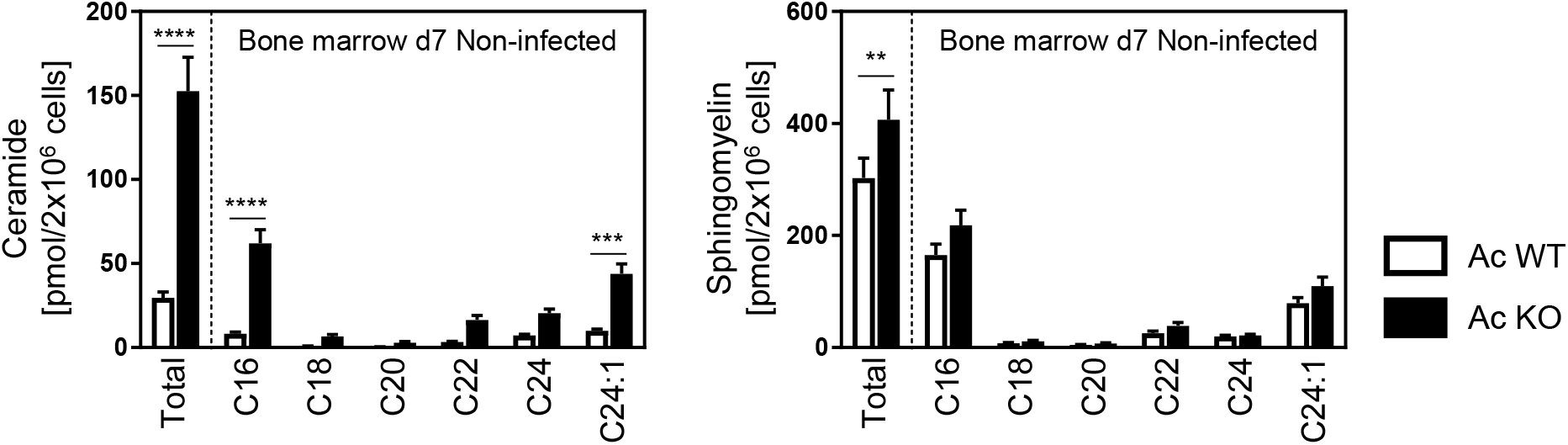
Sphingolipid level of non-infected Ac KO and Ac WT mice. Ceramide (left panel) and sphingomyelin (right panel) levels in bone marrow of non-infected Ac WT and Ac KO on day 7 were determined by HPLC-MS/MS (n=7-8). Data are presented as mean ± SEM. Statistical analyses were performed using two-way ANOVA, followed by Sidak’s post-test (\*\**p*<0.01, \*\*\**p*<0.001, \*\*\*\**p*<0.0001).

